# Neurobehavioral Correlates of How Time-on-task and Sleep Deprivation Modulate Deployment of Cognitive Effort

**DOI:** 10.1101/865766

**Authors:** Stijn A.A. Massar, Julian Lim, Karen Sasmita, Bindiya L. Ragunath, Michael W.L. Chee

## Abstract

Sustaining attention is highly demanding and can falter if there is a shift in willingness to exert effort. Motivated attentional performance and effort preference were tracked in relation to increasing time-on-task (Experiment 1) and sleep deprivation (Experiment 2). Performance decrement with time-on-task was attenuated with reward, while preference to deploy effort decreased with longer task duration. Sleep deprivation, accentuated performance decline with time-on-task, and was accompanied by greater effort-discounting. Motivated attention performance was associated with higher fronto-parietal activation, in both normal and sleep deprived conditions. However, after sleep deprivation modulation of activation by reward was reduced in the anterior cingulate cortex (ACC) and left anterior insula (aIns). Together, these results depict how motivational decline affects performance when one gets tired after sustained task performance and/or sleep deprivation.

## Introduction

Prolonged performance of an attention demanding task results in slowing of responses and increased errors (Warm, Parasuraman, Matthews 2008) and constitute time-on-task effects. Accompanying these behavioral alterations is reduced fronto-parietal activation (Coull et al., 1998, Warm and Parasuraman, 2007, Langner and Eickhoff, 2013, Lim et al., 2010) that reflects decline in top-down attentional control. Resource theories of fatigue posit these that these phenomena arise because prolonged task performance depletes finite neural resources.

Sustained attention is highly sensitive to sleep deprivation (SD) (Lim and Dinges, 2008). Moreover, time-on-task effects can be exacerbated after SD (Van Dongen et al., 2011). Performance deficit in SD arises from state specific mechanisms such as the stochastic drop-out of neural activity in cortical columns arising from ‘local sleep’(Vyazovskiy et al., 2011) or changes in brain connectivity (Yeo et al., 2015, Wang et al., 2016), However, in common with time-on-task effects, SD is also accompanied reduced activation in fronto-parietal brain networks (Chee and Tan, 2010, Chee et al., 2008, Lim et al., 2007, for a review see Ma et al., 2015). Indeed, brain regions showing reduced activation with time-on-task and SD show significant overlap (Asplund and Chee, 2013).

While resource depletion accounts remain the most common explanation for the observed effects of time-on-task and SD, it has recently been argued that both conditions are accompanied by a loss in motivation to exert appropriate effort (Massar et al., 2019a, Müller and Apps, 2019). Underlying both observations is the theory that the brain continuously compares energetic costs against expected rewards (or benefits), especially when one is tired (Boksem and Tops, 2008). If task costs outweigh the benefits, an individual may decide to withdraw effort and reduce standards of task performance (Hockey, 2013, Kanfer and Ackerman, 1989). Such a shift in effort preference may contribute to the observed reductions in brain activation and performance (Massar et al., 2018, Müller and Apps, 2019).

This cost-benefit analysis is thought to be undertaken by the anterior insula (aIns), dorsolateral Prefrontal Cortex and dorsal portion of the medial prefrontal cortex (dmPFC; Boksem and Tops, 2008, Müller and Apps, 2019). The anterior insula has been advanced as an important hub for evaluating task performance. It processes interoceptive information about the internal state of the organism (Craig, 2009), encodes the subjective perception of effort (Otto et al., 2014), and signals if it is appropriate to cease effortful action (Meyniel et al., 2013). Complementing these functions, the dorsal medial prefrontal cortex (dmPFC), including the anterior cingulate cortex (ACC) integrate information about expected rewards and effort costs (Klein-Flügge et al., 2016, Prevost et al., 2010, Shenhav et al., 2013, Vassena et al., 2017). Moreover, the ACC may direct motor and cognitive areas to prioritize high value processes. The lateral Prefrontal cortex tracks subjective cognitive effort, biasing action away from high effort task options (McGuire and Botvinick, 2010). Together, these brain areas are thought to implement the proposed decision process by integrating cost and benefit information given the internal state, computing subjective values of actions, and then energizing the action chosen on the basis of its highest subjective value (Chong et al., 2016, Pessiglione et al., 2017).

In this study we examined the motivational account of performance decrement under two related yet distinct contexts where it is expected to operate i.e. time-on-task (Experiment 1) and sleep deprivation (Experiment 2). Participants underwent fMRI while performing an effortful attentional task under different reward conditions (Figure 1A). Changes in performance and brain activation were tracked for the different incentive levels. To examine the underlying basis for motivated behavior, participants performed an out-of-scanner value-based decision task (discounting task; Figure 1B), in which they pitted the value of potential rewards against the costs of performing the attention task for a specified duration. We showed that attentional performance deteriorated with time-on-task, and that this effect was less pronounced under the high incentive condition. In the decision task, participants discounted the value of available rewards when longer task performance was required. Moreover, both deterioration of attentional performance and value discounting were accentuated after sleep deprivation. Lastly, improved performance with higher reward was associated with greater activation of attention-related brain areas including the aIns, and ACC/dmPFC. Conversely, activity in these areas was reduced with time-on-task and sleep deprivation.

**Figure 1.**
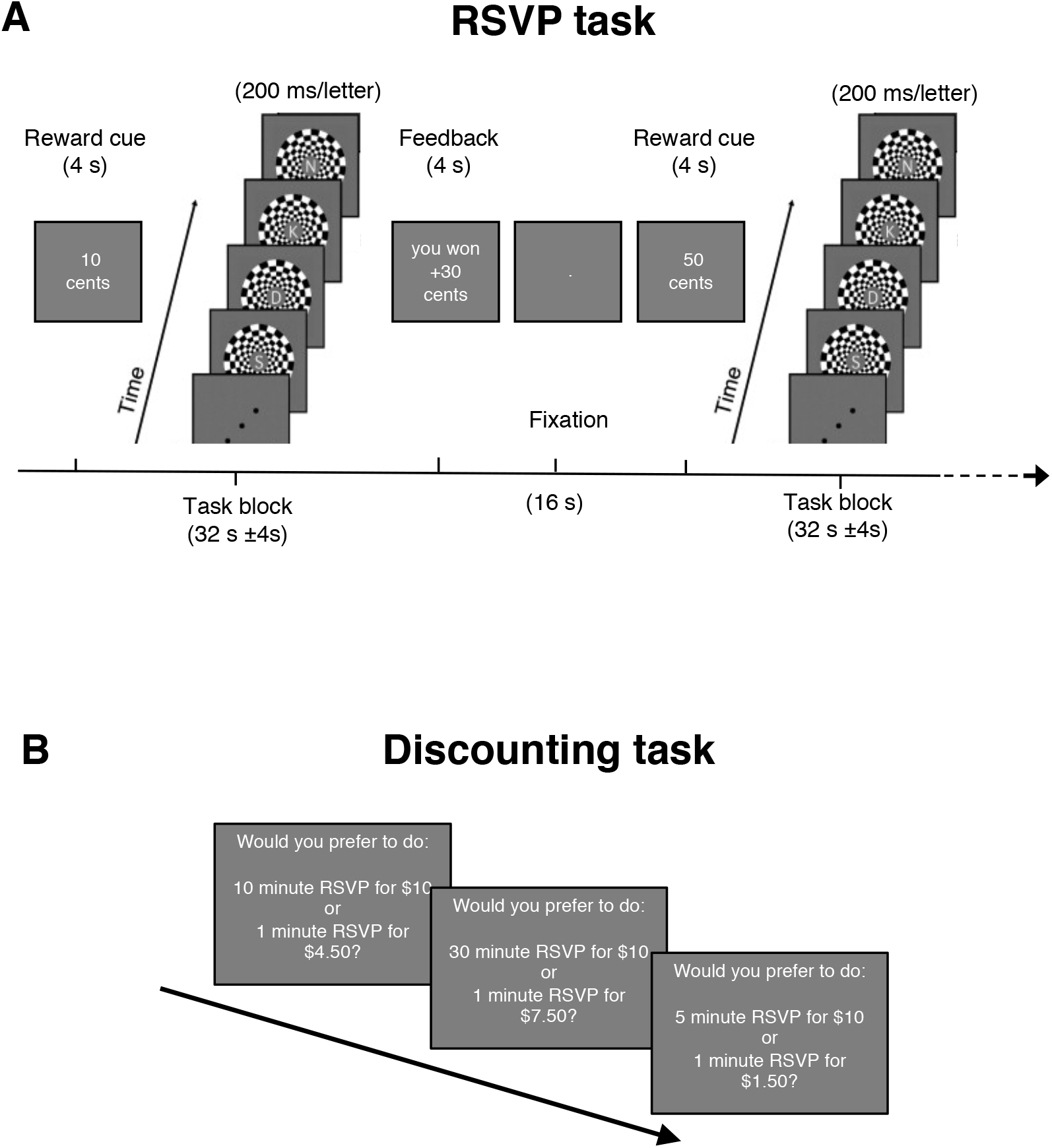
Schematic of experimental tasks, with (A) the rapid serial visual processing (RSVP) task. Letter streams (200ms/letter) are presented in 32 sec (± 4sec) task blocks. Participants had to detect “J” or “K” target letters and respond with a left or right button press, respectively. Each correct target detection was rewarded. Task blocks were preceded by a reward cue, that indicated the incentive level for correct detections in that block (0, 10, or 50 cent). Reward feedback was given after each block. Each task run consisted of six task blocks. A total of six task runs were performed. (B) Example choice trials in the discounting task, in which further performance of the RSVP task (1 to 30 minutes) was weighed against available reward ($0 to $10). After completion of the discounting task, one choice trial was randomly drawn for execution.

## Results

### Motivated Attention Performance

Participants performed six runs of a rapid serial visual processing task (RSVP). Each run consisted of six task blocks during which participants viewed a rapid stream of letter stimuli. Participants had to detect the target letters “J” and “K” and responded by pressing a left or right button, respectively. Each block was preceded by a reward cue that signaled the level of reward (low = 0 cent, medium = 10 cents or high = 50 cents) that could be earned for correct and fast target responses.

#### Experiment 1

Detection accuracy for the RSVP task (Fig. 2A) was analyzed using a repeated measures ANOVA with Reward (low, medium, high), and Time-on-task (run: 1, 2, 3, 4, 5, 6) as within-subject factors. There was a significant main effect of Reward (F(2, 46) = 5.98, p = .015, η_p_^2^ = .206), but no main effect of Time-on-task (F(5, 115) = .955, p = .45) or Reward x Time-on-task interaction (F(10, 230) = 1.77, p = .068, η_p_^2^ = .071). However, a linear interaction contrast (taking into account the first and last runs) was significant (F(1, 23) = 5.017, p = .035, η_p_^2^ = .179). Planned t-tests showed that the time-on-task decrement was significant only in low reward task blocks (t(23) = 2.06, p = .05), but not for task blocks with medium or high rewards (medium: t(23) = −.49, p = .64; high: t(23) = −1.16, p = .26).

**Figure 2.**
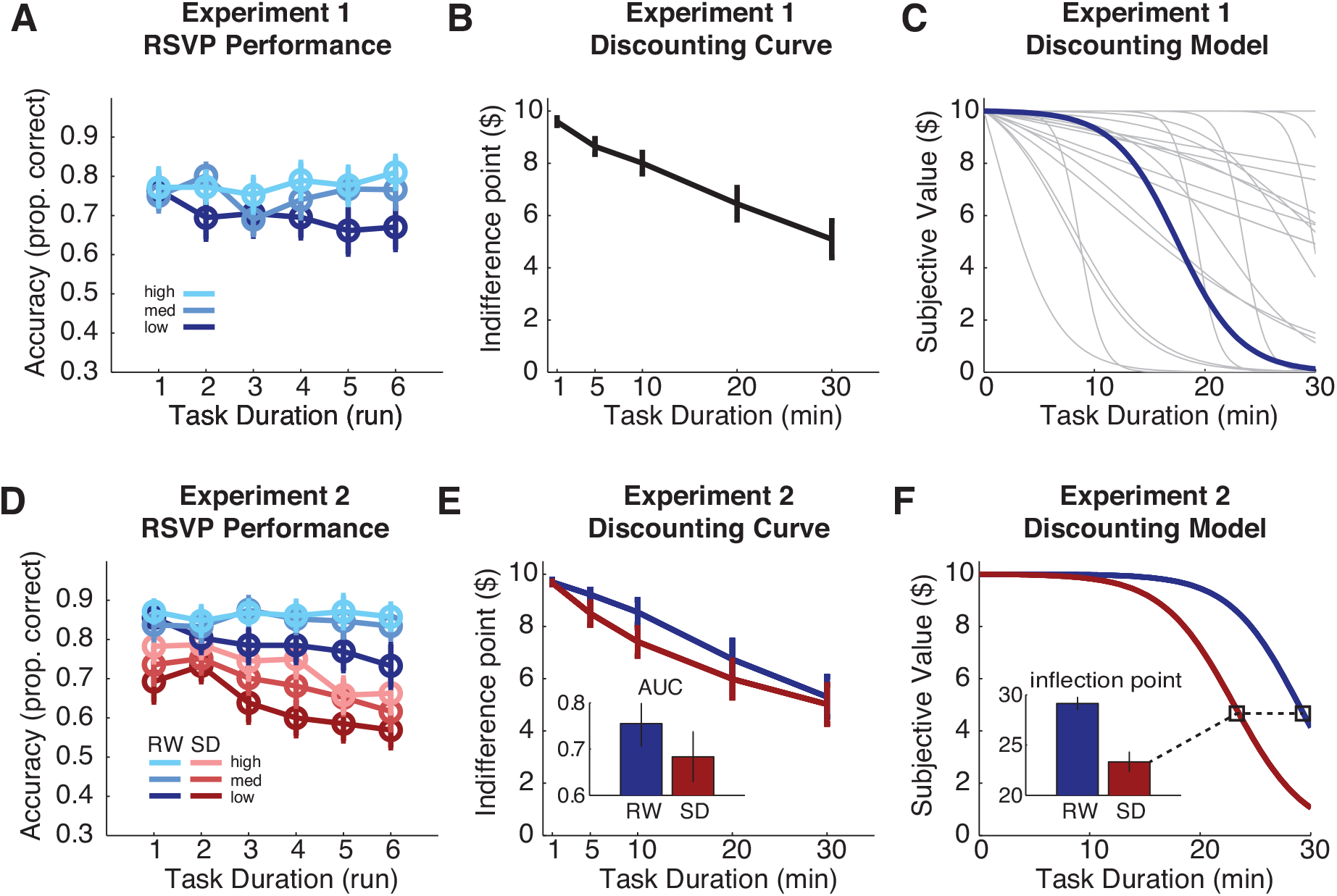
Behavioral results for Experiment 1 (upper panels), and Experiment 2 (lower panels) showing performance on RSVP task at different levels of reward (A & D), Model-free analysis of choice preference in the Discounting task (B & E: with inset in panel E showing area under the curve [AUC]), and computational model fit of the Discounting task (C: light grey lines show individual fits; F: inset shows inflection point of discounting model [p-parameter]). RW = Rested Wakefulness, SD = Sleep Deprivation.

#### Experiment 2

Participants performed the same motivated RSVP task once after a night of normal sleep (Rested Wakefulness: RW), and once after a night of total sleep deprivation (SD). Accuracy data for Experiment 2 are displayed in Fig. 2D. Analysis of the RW session mostly replicated results from Experiment 1. There was a significant main effect of Reward (F(2, 50) = 8.71, p = .002, η_p_^2^ = .258), but no main effect of time-on-task or interaction. This time, the linear interaction contrast was not significant (F(1, 25) = 3.565, p = .071, η_p_^2^ = .125).

To examine the changes in accuracy under SD, a State (RW, SD) x Reward (low, medium, high) x Time-on-task (run: 1, 2, 3, 4, 5, 6) repeated measures ANOVA was performed. There were significant main effects of State (F(1, 25) = 47.21, p < .001, η_p_^2^ = .654), Reward (F(2, 50) = 16.05, p < .001, η_p_^2^ = .391) and Time-on-task (F(5, 125) = 5.341, p < .001, η_p_^2^ = .186) as well as a significant State x Time-on-task interaction (F(5, 125) = 3.059, p = .012, η_p_^2^ = .109), indicating that overall the time-on-task decrement was more pronounced in SD than RW. All other interaction effects were non-significant (all F’s < 1.02).

### Discounting Task

Following completion of the RSVP task, participants performed a value-based decision task in which they were offered monetary rewards for performing the RSVP task for an extended duration (1 to 30 minutes). On each trial of the discounting task participants were presented with a choice between performing the RSVP for a short duration to receive a small reward (<$10), or performing the RSVP for a longer duration to receive a larger reward ($10). Participants had to indicate their preference on each trial. Reward values were systematically updated to approach the participant’s individual indifference point, reflecting the discounted reward value that the participant considered equally attractive as $10 at a longer task duration (subjective value).

#### Experiment 1

Analysis of the discounting curve showed that reward value was significantly discounted with longer task duration (F(1, 23) = 44.91, p < .001, η_p_^2^ = .661; Fig2B). While the value of a $10 reward was discounted slightly when a 5-minute RSVP was required (mean indifference point = $8.64 (1.46)), it was discounted more strongly when 30 minutes of RSVP performance were required (mean indifference point = 5.09 (3.45)). Computational modeling (see Methods) of the choice data indicated that the Sigmoid discounting model (Fig 2C) fit the data better than other models (Hyperbolic or Exponential). This concurs with recent findings concerning effort-based decision making (Klein-Flügge et al., 2015, Massar et al., 2019b). The sigmoid discounting model is characterized by three free parameters, 1) an inflection point (*p*-parameter; indicating the task duration at which the offered reward is discounted to half its original value), 2) a slope (*k*-parameter; indicating the steepness of discounting around the inflection point), and 3) an inverse temperature parameter (Softmax *β* parameter), indicating how strongly choices were determined by the given value function (inverse randomness).

#### Experiment 2

In Experiment 2 reward value was similarly discounted with longer task durations (Duration main-effect: F(4, 100) = 26.98, p < .001, η_p_^2^ = .519). Further there was a significant State main-effect (F(1, 25) = 5.44, p = .028, η_p_^2^ = .179), showing that reward value was more heavily discounted under SD compared to RW. The state effect was further examined through two additional methods. First, we extracted the area under the discounting curve (AUC), which is a model-free measure of discounting (Myerson et al., 2001). A paired t-test showed that the AUC was significantly lower in SD compared to RW (t(25) = 2.32, p = .029, Cohen’s *d* = .455; Fig. 2E inset). Secondly, we fitted a computational model to the individual choice data. Comparison of the individual model parameters between SD and RW showed that the Sigmoid slope (k-parameter) was not altered between SD and RW (t(25) = 1.33, p = .195). However, there was a significant shift in the inflection point (p-parameter), such that in SD compared to RW, shorter durations of required task performance led to discounting of a reward to half its original value (t(25) = 2.09, p = .047, Cohen’s *d* = .41; Figure 2F). Furthermore, choice randomness was higher (i.e. Softmax *β* parameter was lower) in the SD session compared to the RW session (t(25) = 2.204, p = .037, Cohen’s *d* = .43).

### Imaging results

To examine BOLD activation during performance of the motivated RSVP task, a general linear model (GLM) analysis was performed, including a regressor modeling the overall activation during task blocks. An additional Reward regressor modeled the parametric modulation of task activation with reward level (indicating increasing activation during higher reward blocks).

#### Experiment 1

Analysis of the main task regressor (Fig 3A) showed large clusters of activation in bilateral occipital cortex, as well as activation in attention related areas comprising prefrontal clusters in the bilateral inferior frontal gyrus stretching to the anterior insula (aIns), bilateral superior frontal gyrus stretching to the cingulate gyrus, and lateral prefrontal clusters comprising the left precentral and middle frontal gyrus, and right inferior frontal gyrus, and parietal clusters in the right precuneus and left superior parietal lobule. Further, both caudate nuclei were activated.

**Figure 3.**
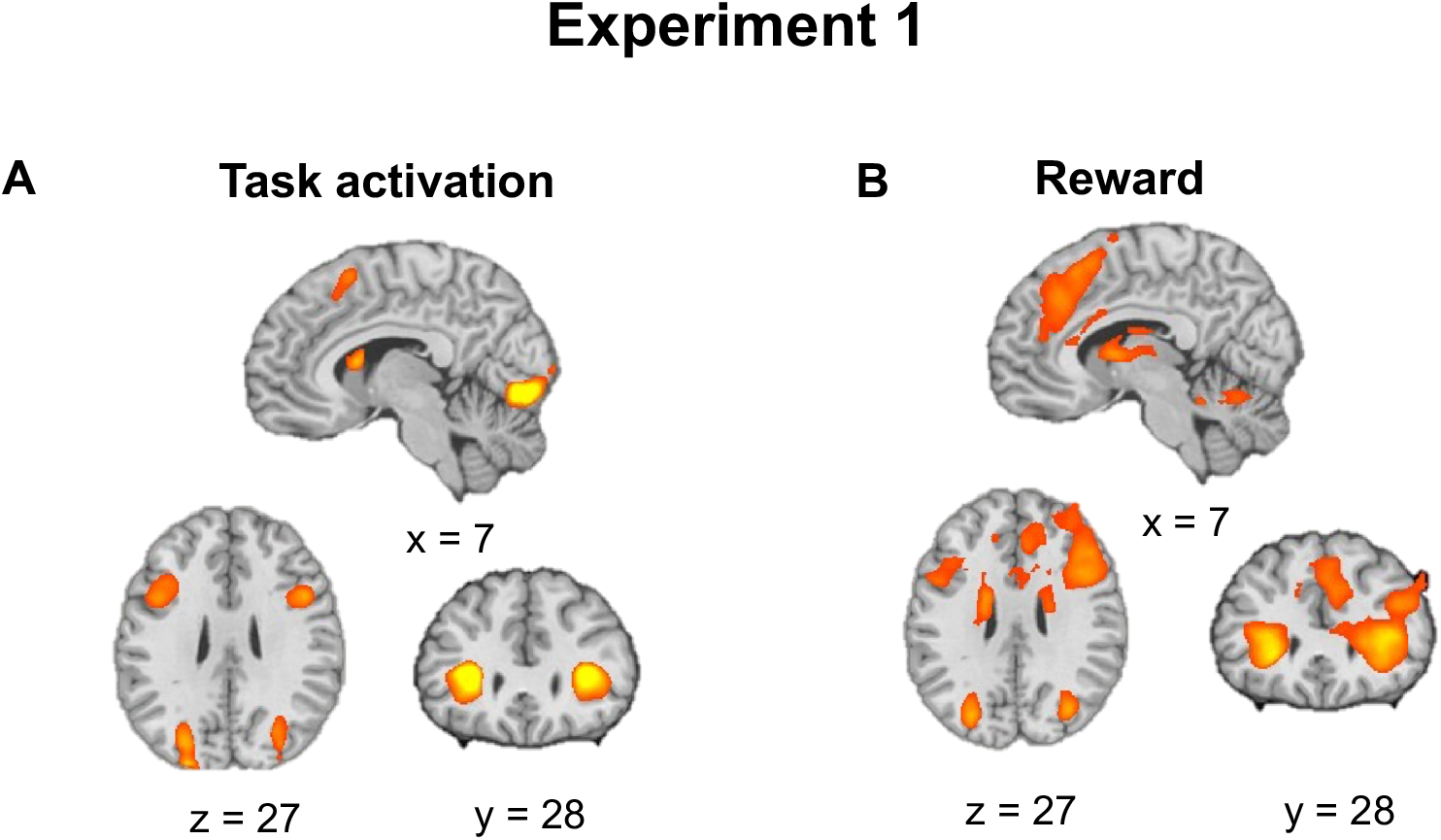
Activation during the RSVP task in Experiment 1. A) main effect of task, B) parametric modulation of activation by reward.

#### Reward modulation

Activation in areas engaged during task performance increased with incentive value. Moreover, clusters that were modulated by reward extended medially into the anterior cingulate cortex (ACC), and laterally into the superior and middle frontal gyrus, mainly right lateralized. Parietal activation was also modulated by reward, involving the inferior parietal lobule bilaterally. Several additional areas showed reward modulation including clusters in the ventral striatum, thalamus and cerebellum.

#### Time-on-Task

To examine changes in activation over time-on-task, a second GLM analysis was performed, modeling task activation in the six task runs separately (with separate regressors for each reward level, in each run). Systematic changes in activation over time (indicated by a significant run main effect) were found in visual areas, and in the medial frontal gyrus/cingulate cortex, middle frontal gyrus, and precuneus (Figure 4A). Despite activation declining over time-on-task in all three reward conditions, high reward trials continued to show greater activation that low reward trials over time (Figure 4B).

**Figure 4.**
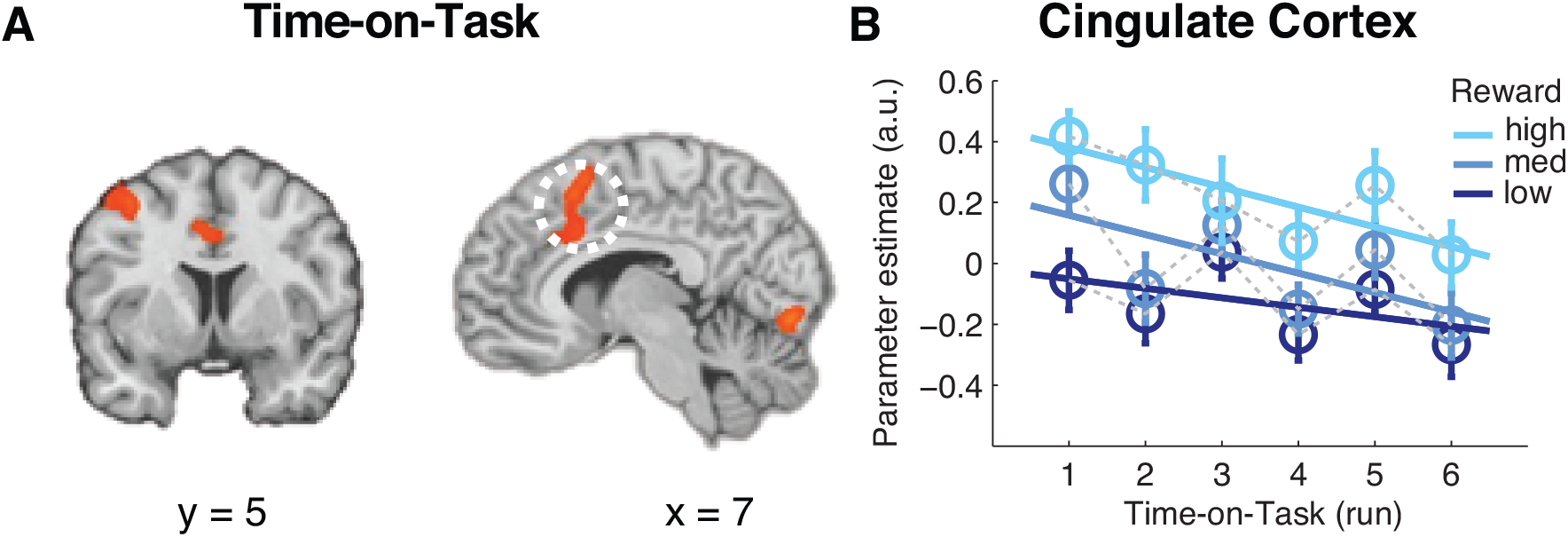
A) Time-on-Task effect in task activation, B) time course of task activation in dmPFC/ACC.

#### Experiment 2

The main effect of task seen in Experiment 1 was also present in both the RW and SD sessions of Experiment 2 with activation of visual cortex, and fronto-parietal areas Fig 5A). In RW, activation in the attention-related areas was up-regulated in higher reward blocks with extension to additional areas of cingulate cortex, striatum and the thalamus (Fig 5B). In SD this reward modulation appeared in similar brain regions, but was weaker (Fig 5B). Direct contrast of the reward modulation during RW versus SD yielded two clusters, a significant cluster in the left anterior insula and a small patch of medial frontal gyrus, that did not survive cluster level correction (see Fig 6).

**Figure 5.**
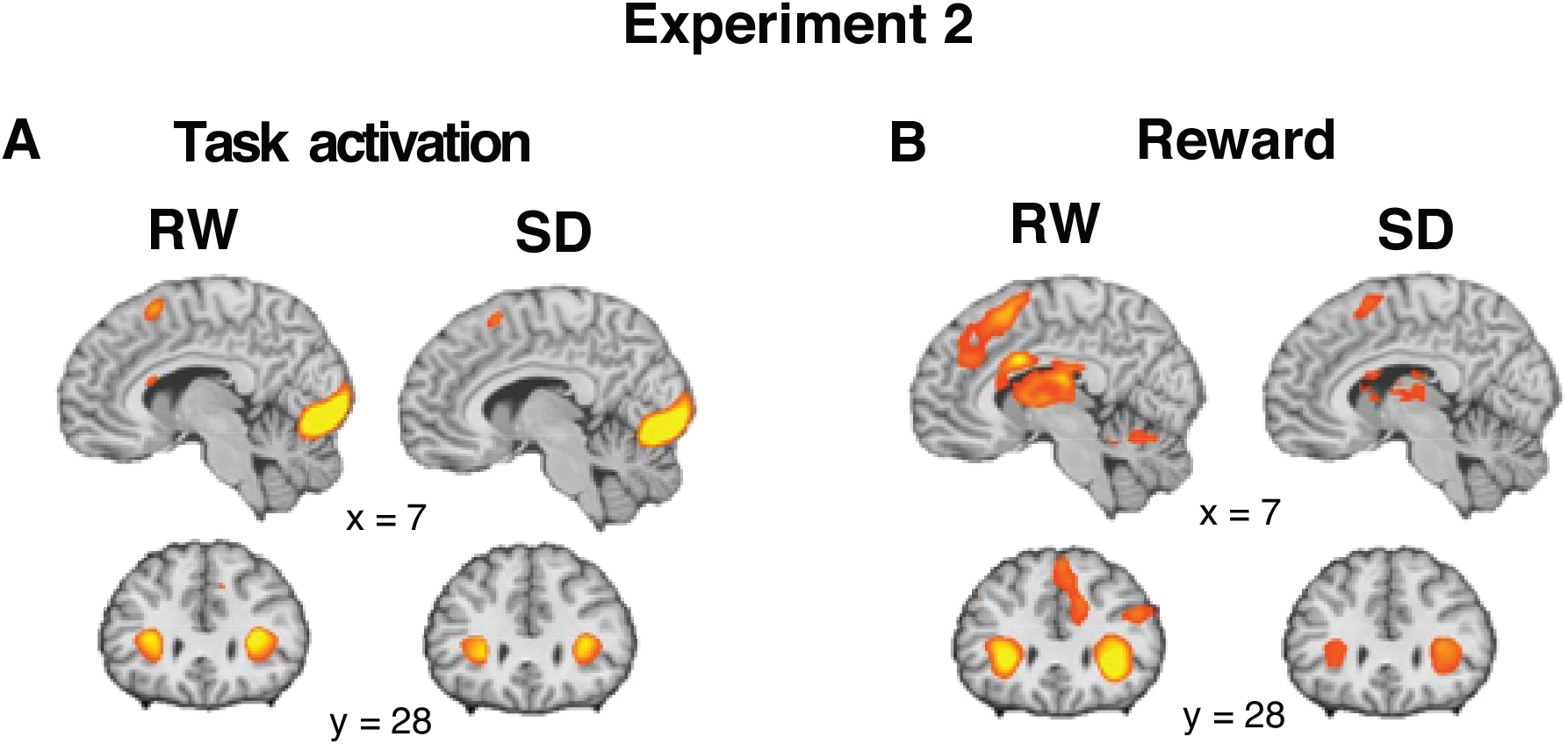
A) Task activation in Experiment 2 during Rested Wakefulness (RW) and Sleep Deprivation (SD). B) Parametric modulation by reward in RW and SD.

**Figure 6.**
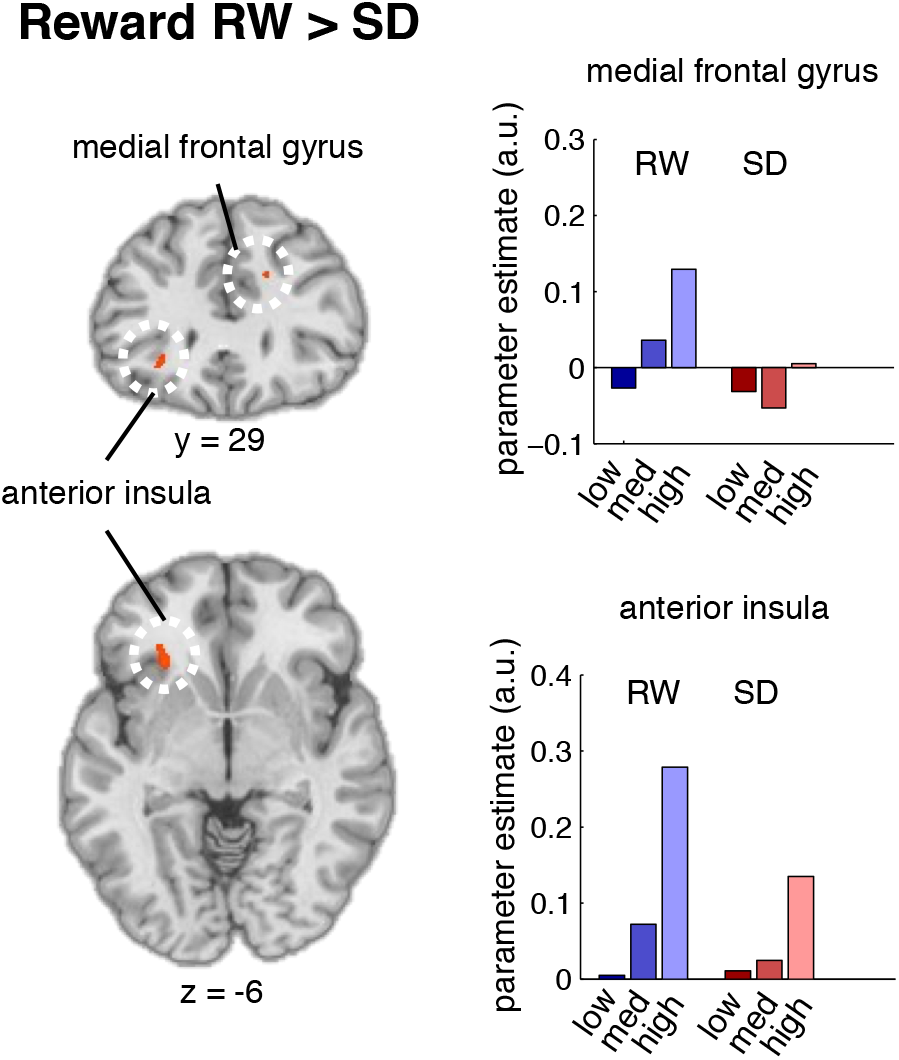
Areas showing reduction in reward modulation during SD compared to RW with, medial prefrontal cortex (upper panels), and left anterior insula (lower panels). Bar graphs show task activation in the indicated medial frontal gyrus and anterior insula clusters, modeled for each reward condition.

## Discussion

We studied the effects of motivation, time-on-task and sleep deprivation on task performance as well as their interactions. We also measured discounting behavior, and brain activation associated with task performance under different conditions. In addition to replicating previous findings that task duration and sleep deprivation negatively impact the behavioral and neural correlates of motivated behavior, we found that: 1) Performance was influenced by reward motivation and by sleep deprivation, 2) Reward value was discounted with longer task durations and with SD, 3) Brain activation in areas mediating attentional control was lowered with time-on-task but tempered by reward particularly during SD. Overall, these data underscore the interaction between effortful task performance and motivation, and support a motivational account of fatigue and its modulation by sleep deprivation.

### Performance decrement is modulated by motivation

Deterioration of attentional performance over time is one of the most robust behavioral hallmarks of prolonged task performance (for reviews see Langner and Eickhoff, 2013, Warm et al., 2008). The time-on-task decrement is often magnified by task load as well as by sleep deprivation (Lim and Dinges, 2008). This is commonly taken to reflect that the participant is no longer able to muster the energetic resources to maintain performance. Results from the current study show that performance declines over time can be mitigated by the provision of incentives. In conditions of normal sleep (Exp1 & Exp2 RW), performance only deteriorated in the low reward blocks, while high detection accuracy was maintained throughout in the higher reward blocks. This is in line with the idea that when fatigued, resources can still be directed to facilitate task performance if the reward for maintaining performance is sufficiently high. Several earlier studies have shown similar reward modulation of time-on-task effects on shorter time scales (i.e. ~10 minutes; Esterman et al., 2016, Massar et al., 2016). Other studies have shown that, even after sustained task performance for 90 minutes or longer, performance can be boosted if incentives for performing well are provided (Boksem et al., 2006, Hopstaken et al., 2014, but see Gergelyfi et al., 2015).

As expected, performance was impaired with sleep deprivation compared to rested wakefulness in Experiment 2. Furthermore, time-on-task related performance decline was steeper during SD. While reward motivation modulated performance at all points in time, it did not interact with the time-on-task effects in SD. Although this only partially replicated results from an earlier study (Massar et al., 2019b), attention could still be substantially improved by high reward during SD. Of note, the worst performance in the high reward blocks (run 6) was at the level of low-reward performance in the first run (around 60% accuracy).

### Reward value is discounted with task duration

The discounting task provided further insight into the proposed decision processes that may underlie the withdrawal of attentional effort with longer task duration. Participants were given the free choice between performing the attention task for an additional short period of time (1 minute) and receiving a smaller reward, or performing the task for a specified longer duration (up to 30 minutes) for a higher reward. When longer task durations were required, participants often opted to forego the higher reward by choosing a shorter task.

These findings demonstrate that task performance is considered as a cost by which reward value is discounted (Westbrook and Braver, 2015). Choice behavior best fit a sigmoid discounting function where with small increases in effort, reward value is minimally discounted, while with larger increases in effort, discounting becomes steeper (i.e. the discounting function is initially concave). Similar discounting functions were found when effort was manipulated along a dimension other than task duration (e.g. cognitive task difficulty or physical exertion; Chong et al., 2017, Klein-Flügge et al., 2016, Klein-Flügge et al., 2015, Białaszek et al., 2017). Importantly, discounting was more pronounced after sleep deprivation, paralleling the finding that performance decline with time-on-task in the RSVP task was exacerbated during SD. The observation that task duration had a stronger influence on preference in SD than in RW concurs with the finding that effort discounting also increases after SD (Libedinsky et al., 2013, Massar et al., 2019b) and suggests a loss in motivation to perform.

### Motivated performance is supported by a cingulate-insula network

A third finding from the current study was that reward modulates activation of brain areas involved in top-down control of attention. A network of frontal (aIns, dmPFC, lateral PFC) and parietal areas (precuneus/IPL) that was involved in overall task performance, was modulated by incentive value. Under normal sleep conditions (Exp1 & Exp2 RW) reward modulated activation extended into the ACC, and lateral middle and superior frontal gyrus, and into clusters in the thalamus and striatum. With time-on-task, activation in the ACC/dmPFC, middle frontal gyrus and precuneus decreased. These areas closely match the regions found in a previous fMRI study (Asplund and Chee, 2013). The modulation of aIns activation by reward decreased following SD.

The insula, ACC/dmPFC and lateral frontal cortex are thought to be key nodes in a network integrating reward value and effort costs (Vassena et al., 2017), and coordinating the allocation of effort to courses of action with the highest net gain (Pessiglione et al., 2017). In several meta-analyses, these areas show remarkable overlap in their response to reward for task performance (Parro et al., 2018), time-on-task (Langner and Eickhoff, 2013), and sleep deprivation (Ma et al., 2015, for a comparison of overlap see Massar et al., 2019a). As such this network may serve to redirect neural resources when effort is perceived to outweigh rewards when one is tired (Müller and Apps, 2019), or in our case, sleep deprived.

These areas have repeatedly featured in effort-based decision making tasks-involving exertion of physical force (Bonnelle et al., 2016, Burke et al., 2013, Klein-Flügge et al., 2016, Prevost et al., 2010), and cognitive effort (Chong et al., 2017, Massar et al., 2015). Activation in the ACC during effort-based decision making is correlated with individual differences in self-reported persistence (Kurniawan et al., 2010). Interestingly, electrical or magnetic stimulation of the medial frontal cortex can induce a sense of motivation (Parvizi et al., 2013), and bias choices to higher effort options (Zenon et al., 2015). The current data add to these findings that both time-on-task and sleep deprivation reduce activation in this network, potentially reflecting the reduced motivation to exert effortful control.

### Conclusion

In summary, the current study shows that motivation and attention interact through a fronto-parietal brain network. With increasing time-on-task and sleep deprivation, performance and willingness to exert effort deteriorate. This is accompanied by reduced activation in the ACC and the aIns, suggesting a key-role for these areas in integrating the costs and benefits of cognitively effortful performance, and the modulation of how these costs and benefits are perceived when we get tired.

## Methods

### Participants & Procedure

#### Experiment 1

Twenty-four participants were recruited from the student population of the National University of Singapore (10 females, mean age (stdev.) = 22.7 years (2.26)). All participants were right-handed, had no contra-indications for MRI scanning, and had no history of psychiatric or neurological disorders. The protocol was approved by the Institutional review board of the National University Singapore, and all participants provided informed consent prior to testing. Participants came to the lab for one session, during which they were scanned while performing a motivated attention task. This was followed by an out-of-scanner value-based decision-making task (Discounting). This session lasted approximately 1.5 hours. Participants were paid $25 for their time, plus a performance-dependent bonus of up to $25.

#### Experiment 2

An independent sample of 28 participants was recruited for a sleep deprivation experiment (15 females, mean age (stdev.) = 22.9 years (3.28)). The same inclusion criteria as for Experiment 1 were applied, with additional criteria that participants should have regular sleeping habits, no symptoms or history of sleep disorders, and should not work irregular or night shifts. Participants in Experiment 2 were studied in two sessions. During the Rested Wakefulness (RW) session they slept in the lab (bedtime 11pm to 7am). They were then woken up and the experimental session started at 8am. In the Sleep Deprivation (SD) session participants were kept awake overnight, supervised by a research assistant. They were allowed to engage in non-strenuous activities. In the morning, the testing session commenced at 6am. This is when cognition is usually most strongly affected by the combination of the circadian factors and the effects of extended wakefulness. In both the RW and SD sessions testing procedures were the same as in Experiment 1. Participants performed a motivated attention task, and a Discounting task. Participants received $80 for completion of both sessions, plus a performance-dependent bonus of up to $25 per session.

### Motivated Attention Task

Participants performed an attention-demanding task (Asplund and Chee, 2013) while in the scanner. A rapid serial visual presentation (RSVP) stream of white letters was presented on a grey background, surrounded by a flickering checkerboard (10 Hz). Letters were presented for 200 ms each, in direct succession. Participants were required to press one of two target buttons whenever they detected a “J” or a “K” letter in the RSVP stream. Task blocks lasted for 32 ± 4 seconds. Each task block contained 5 to 7 target letters separated by at 2 to 10-seconds inter target interval. Critically, each task block was preceded by reward cue (1c, 10c, or 50c), that informed participants of the incentive that could be earned for correct and fast target responses during that block. An RT cut-off was determined based on the median RT of an out-of-scanner practice run. A task run lasted for 6.5 minutes and comprised 6 task blocks (2 blocks per incentive level in counterbalanced order). Participants performed a total of 6 task runs. The main performance metric was target detection accuracy, which was quantified per run for each incentive level separately. Prior to scanning, participants performed a practice run outside the scanner.

### Discounting Task

After completion of the motivated RSVP Task, participants performed an out-of-scanner Discounting Task. In this task participants were to decide how they would spend the last 30 minutes of the experimental session, indicating their preference between performing the RSVP task for another duration of time, given a specified reward. Participants were presented with a series of choice trials. On each trial they could choose to perform a short version of the RSVP task (1 minute) and receive a low reward, or a longer version of the RSVP task (5, 10, 20, or 30 minutes) to receive a higher reward. The higher reward for the longer duration task was always $10. The lower reward was systematically varied from trial to trial using an adaptive staircase method. If the participant chose to do the longer task, the lower reward for the shorter task was adjusted upwards in the next trial. If the participant chose to do the shorter task, the reward amount for the shorter task was adjusted downwards for the next trial. This procedure allowed the estimation of the lower amount of money an individual considered equally valuable as the higher amount $10 at a given task duration (indifference point). A discounting curve was constructed by plotting the indifference points at all different task durations, from which the area under the curve (AUC) was calculated as a model-free summary metric of the individual’s extent of discounting (larger AUC indicates less discounting). To make choices incentive compatible, one choice trial was drawn at random for execution after all choices were made. Participants had to perform the RSVP task for the duration of the chosen option on that trial, before receiving the associated reward. To ensure that decisions were not made on the basis of the delay to reward receipt (delay discounting), all participants had to stay in the lab for a fixed duration of 30 minutes.

#### Model-based analysis

To more formally characterize the shape of the discounting curve, we fitted a sigmoid function previously found to describe effort-based choice well (Klein-Flügge et al., 2016, Klein-Flügge et al., 2015, Massar et al., 2019b). This model predicts that a reward is discounted by small amounts when only a short duration of task performance is required. At longer task durations, discounting becomes steeper and eventually plateaus to approach zero at very long durations of task performance (see Fig7 A&B).

**Figure 7.**
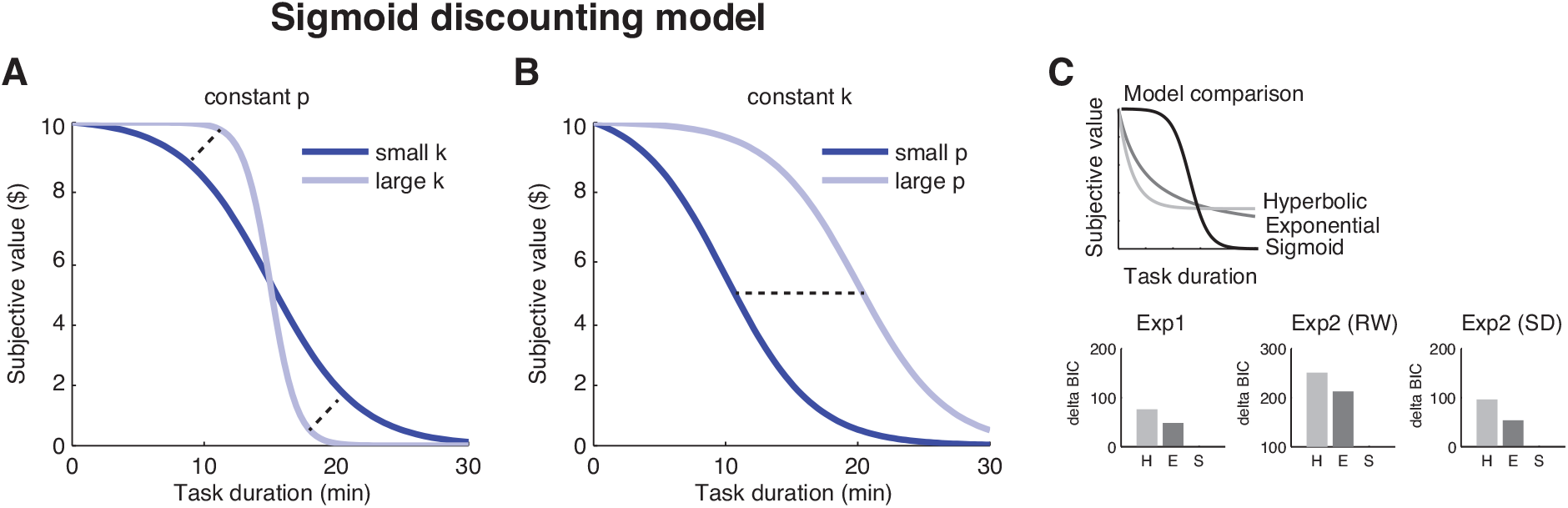
Illustration of the sigmoid discounting model with (A) different values of k (given constant p), (B) different values of p (given constant k), and (C) comparison of model fit against hyperbolic and exponential models. BIC = Bayesian Information Criterion, H = Hyperbolic, E = Exponential, S = Sigmoid.

This model can be formalized by the following equation:

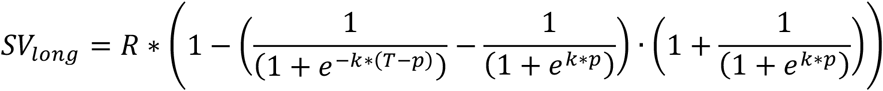

Where *SV_long_* is the subjective value of a reward *R* at task duration *T*. The model has two free parameters that are fitted to individual choice behavior. The parameter *k* indicates the slope of the discounting function, while *p* indicates the inflection point (time point at which the reward is discounted to half its original value). Choice probability for each trial was modeled following a Softmax function:

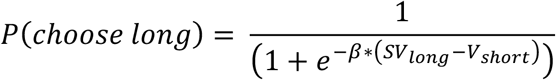

Where *P* is the probability of choosing the longer task. The free parameter *β* denotes the inverse choice randomness (inverse temperature), indicating how strongly choices follow the given value function (i.e. the influence of value difference between the subjective value of the longer task option *SV_long_*, and the value of the shorter duration option *V_short_*). Individual participants’ choice data were fitted using the fmincon algorithm in Matlab (MathWorks, Natick, MA) to find free parameter values that optimized the model fit (minimize negative log likelihood). To test whether the sigmoid model described choice data well, model fit was compared to more traditional hyperbolic and exponential discounting models using the Bayesian information criterion (BIC; Schwartz, 1978). A smaller BIC indicates better model fit (Fig7 C, where delta BIC indicates the difference score with regard to the best fitting model). To compare model parameters between the RW and SD session in Experiment 2, paired-samples t-tests were performed for the sigmoid *p*, *k*, *β*-parameters (square root transformed to correct for non-normality; Peters et al., 2012).

### fMRI data acquisition & processing

Functional imaging during the Motivated Attention Task was conducted on a Siemens 3T Prisma scanner (Siemens, Erlangen, Germany). Functional runs were acquired using an interleaved echo-planar imaging (EPI) sequence (TR: 2000ms; TE: 30ms; flip angle: 90°; field-of-view: 192 × 192 mm; matrix size: 64 × 64). Thirty six 3-mm oblique axial slices were collected, aligned to the intercommissural plane. A T1-weighted high-resolution 3D-MPRAGE (1mm × 1mm × 1mm) sequence (TR: 2300 ms; TE: 2.28 ms; TI = 900 ms; flip angle = 8°; field-of-view: 256 × 240 mm; BW = 240 Hz/Px; matrix size: 256 × 240, voxel size: 1 mm^3^; 192 slices) was performed at the end of the functional runs. During functional imaging, an MRC 12M-I eye-tracking camera (MRC Systems GmbH, Germany) was used to monitor eye closure. Audio messages were delivered through an intercom if a participant closed their eyes for ≥10 seconds, to reduce episodes of sleep.

#### Preprocessing

Functional imaging data were slice-time corrected and motion corrected using rigid body translation and rotation parameters. Individual participants’ anatomical scans were then reconstructed into surface representations and functional data were registered to structural images using the reconstructed cortical surfaces (Greve and Fischl, 2009, http://surfer.nmr.mgh.harvard.edu/fswiki/FsFast). The structural images were in turn nonlinearly registered to the MNI152 space (Buckner et al., 2011, Yeo et al., 2011). The resulting nonlinear deformations were used to warp the functional data into MNI152 space and smoothed with a 6 mm FWHM smoothing kernel. The first four volumes of each scan run were discarded to allow for signal saturation.

#### Statistical analysis

To analyze overall activation during task performance and the modulation of activation by rewards a GLM analysis with two predictors of interest was performed in BrainVoyager QX version 2.6.1.2318 (Brain Innovation, Maastricht, the Netherlands). The first regressor modeled the overall activation during task performance by modeling task blocks as boxcar functions with the length of the block duration. An additional parametric regressor modeled the incentive level for each block, orthogonalized with respect to the main task regressor ([−1 0 1] for low, medium and high reward). To analyze time-on-task changes in activation, a second GLM with 18 predictors of interest was performed in which task blocks for each reward condition (low, medium, high) were modeled for each task run separately (run 1-6). Areas that showed a significant main effect of task run were further examined to determine the direction of this effect. In all GLMs an additional regressor of non-interest was included to model motor responses. 6 regressors were included to account for head motion. All regressors were convolved with a hemodynamic response function and GLMs were accordingly computed. Resulting statistical maps were thresholded at p < .001 (uncorrected), and corrected (p < .05) using an iterative cluster size thresholding algorithm (Goebel et al., 2006). In Experiment 1, one participant had excessive head motion (> 1 mm displacement in multiple runs), and was excluded from fMRI analysis (final N = 23). In Experiment 2, two participants were excluded from analysis as they were unable to perform the attentional task under SD (final N = 27). One participant completed only five runs of the attention task, and was excluded from the time-on-task analysis (final N = 26). For another participant data from the discounting task were not correctly saved in one session, and were not included in the discounting analysis (final N = 26).

## Acknowledgments

The authors would like to thank James Teng, Teck Boon Teo, and Ksenia Vinogradova for technical assistance and help collecting data. This work was supported by grants awarded to Dr. Michael Chee by the National Medical Research Council (NMRC/STaR/0015/2013) and the Far East Organization.

